# Asymmetric effective connectivity between primate anterior cingulate and lateral prefrontal cortex revealed by electrical microstimulation

**DOI:** 10.1101/369488

**Authors:** Verónica Nácher, Seyed Alireza Hassani, Thilo Womelsdorf

**Author notes:** Corresponding authors: Dr. Verónica Nácher, or Dr. Thilo Womelsdorf.

## Abstract

The anterior cingulate cortex (ACC) and lateral prefrontal cortex (IPFC) of the non-human primate show neural firing correlations and synchronize at theta and beta frequencies during the monitoring and shifting of attention. These functional interactions might be based on synaptic connectivity that is equally efficacious in both directions, but it might be that there are systematic asymmetries in connectivity consistent with reports of more effective inhibition within the ACC than IPFC, or with a preponderance of ACC projections synapsing onto inhibitory neurons in the IPFC. Here, we tested effective ACC-IPFC connectivity in awake monkeys and report systematic asymmetries in the temporal patterning and latencies of effective connectivity as measured using electrical microstimulation. We found that ACC stimulation triggered evoked fields (EFPs) were more likely to be multiphasic in the IPFC than in the reverse direction, with a large proportion of connections showing 2-4 inflection points resembling resonance in the 20-30 Hz beta frequency range. Stimulation of ACC → IPFC resulted, on average, in shorter-latency EFPs than IPFC → ACC. Overall, latencies and connectivity strength varied more than two-fold depending on the precise anterior-to-posterior location of the connections. These findings reveal systematic asymmetries in effective connectivity between ACC and IPFC in the awake non-human primate and document the spatial and temporal patchiness of effective synaptic connections. We speculate that measuring effective connectivity profiles will be essential for understanding how local synaptic efficacy and synaptic connectivity translates into functional neuronal interactions to support adaptive behaviors.

## Introduction

The anterior cingulate cortex and the lateral prefrontal cortex in primates are believed to exchange information during attention demanding tasks and following behavioral feedback (Alexander and Brown, 2011; Shenhav et al., 2016; Rushworth et al., 2011). This information exchange is essential for successful attentional deployment and for adjusting behavioral strategies (Womelsdorf & Everling, 2015; Hayden et al., 2011; Kennerley et al., 2006). Functionally, this information exchange becomes evident in activity correlations during attention shifts and outcome processing in the form of bilateral ACC-IPFC firing correlations (Oemisch et al., 2015), spike-burst to local field potential (LFP) synchronization in the theta, beta, and gamma band (Womelsdorf et al., 2014), burst-triggered LFP power correlations at theta and beta frequencies (Voloh & Womelsdorf, 2017), inter-areal theta-phase to gamma-amplitude correlations (Voloh et al., 2015), and beta and gamma band LFP correlations (Rothe et al., 2011). The synaptic- and circuit-level mechanisms that underlie this heterogeneous pattern of functional correlations are largely unknown. One reason for this knowledge gap is the unknown efficacy of neuronal connections between both brain areas that could indicate how strong, fast, symmetric, and frequency specific neuronal interactions are between both areas. Our study set out to provide this missing link in understanding the interplay of circuits in ACC and IPFC in the awake rhesus monkey.

Previous anatomical studies have shown bidirectional synaptic connectivity between ACC and IPFC (Arikuni et al. 1994; Lu et al., 1994; Morecraft et al., 2012). This bi-directional connectivity might reveal systematic anatomical asymmetries whose functional consequences are poorly understood. For example, quantitative studies have shown more numerous populations of parvalbumin expressing (PV+) inhibitory interneurons in IPFC than in ACC, but larger densities of slower acting calbindin (CB+) expressing interneurons in ACC than IPFC (Dombrowski et al., 2001). These neuron specific differences are likely paralleled by pathway specific differences in the size of boutons and spines (Medalla and Barbas 2010; Medalla et al., 2017), and overall differences in the number of cells that are within the synaptic terminal fields of ACC → IPFC and IPFC → ACC connections (Gabbott and Bacon 1996; Medalla et al., 2017). From a functional perspective, these anatomical anisotropies are expected to be reflected in unique efficacy profiles, and possibly in a unique temporal patterning of neuronal interactions between both areas. However, it is difficult to constrain hypotheses about the precise consequences of specific structural connectivity differences for functional interactions without some ground truth about the efficacy of inter-areal connections.

Our study, therefore, set out to identify the electrical microstimulation parameter range that allows characterizing the strength, timing, temporal patterning and anatomical specificity of ACC → IPFC, and IPFC → ACC connectivity in the awake non-human primate. We found that stimulation evoked fields showed a similar amplitude dependence and multiphasic patterning in both stimulation directions. Beyond these similarities, we revealed prominent differences with shorter IPFC → ACC latencies, and slower temporal beta frequency range resonance following IPFC → ACC stimulation compared to ACC → IPFC stimulation. We believe that these insights provide constraints to understand how structural connections and circuit compositions lead to a unique profile of functional interactions.

## Materials and Methods

### Subjects and surgical procedures

Experiments were conducted in two adult male monkeys (*Macaca Mulatta*) weighing 11 kg (Monkey S) and 8 kg (Monkey H). Both were implanted with a head holder for immobilizing the head during experiments and with a rectangular recording chamber (20 x 28 mm inner dimensions), placed over the right hemisphere, above the IPFC and ACC following stereotaxic coordinates (Paxinos et al., 2008) and magnetic resonance imaging (MRI). All surgical procedures were carried out under general anesthesia and aseptic conditions. The animals were first anesthetized with ketamine (10 mg/kg i.m.), acepromazine (0.5 mg/kg, i.m.) and atropine (0.04 mg/kg, i.m.), followed by isoflurane anesthetic until a surgical level of anesthesia was achieved. Animals were intubated and artificially ventilated with a mixture of oxygen and air. Expiratory CO_2_, electrocardiogram trace and temperature were monitored continuously during surgery. Antibiotics and analgesics were administered after surgery. All procedures followed the guidelines of the Canadian Council of Animal Care policy on the use of laboratory animals and were approved by the Council on Animal Care of York University.

### Electrical stimulation and recording sessions

During the experiments, the monkeys sat in a custom-made primate chair with their head fixed and were kept alert but quiet by receiving drops of juice after each stimulation protocol (controlled by the experimenter). We either stimulated IPFC and recorded single neurons and local field potentials (LFPs) in ACC or vice versa using tungsten microelectrodes (FHC). The stimulation microelectrodes were 250 μm in diameter with an impedance of 0.1 MΩ and the recording microelectrodes were 125 μm in diameter with an impedance of 1.2-2.2 MΩ. Microelectrodes were lowered daily through guide tubes using software-controlled precision microdrives (Neuronitek, ON, Canada). We typically placed two stimulating microelectrodes in one microdrive, using separate adjacent guide tubes mounted to the microdrive (interelectrode distance of 1 mm), for wide bipolar stimulation (Montgomery, 2010). Two recording microelectrodes were placed in a second microdrive in the same manner. The two microdrives were mounted on the recording chamber and the distance between them during the experiment was 6 mm on average. The recorded or stimulated subfields in IPFC (areas 9/46, 46d, 46v) and ACC (areas 32, 24c) (Fig 1a) were identified by projecting each electrode trajectory onto the two-dimensional brain slices obtained from anatomical MRI images, using the open-source MRIcro imaging software (http://www.mricro.com). At the beginning of each experiment, all microelectrodes were connected to a multichannel acquisition processor (Neuralynx Digital Lynx system, Inc., Bozeman, Montana, USA) for data amplification, filtering and acquisition. Spiking activity was obtained following a 600-8000 Hz bandpass filter and further amplification and digitization at a 32 KHz sampling rate. LFP signals were obtained by low-pass filtering at 300 Hz and downsampling to 1 KHz. Microelectrodes in both microdrives were advanced into the brain at the calculated coordinates for the specific IPFC and ACC subfields and until spiking activity was recorded, for making sure that the stimulation microelectrodes were located in gray matter. Bipolar electrical microstimulation proceeded using the StimPulse Electrical Stimulator System (FHC). Constant currents were delivered to the stimulating microelectrodes and consisted of a single pulse or bursts of 2, 4 or 8 pulses each (Fig. 2a). Each pulse was biphasic, rectangular and charge balanced (190 μs width for each phase with 10 μs pulse delay) with the cathodal pulse preceding the anodal. The inter-pulse delay for burst stimulation was 2 ms. During initial experiments, a range of intensities between 20-240 μA with increasing steps of 20 μA, and a range of 300-1000 μA with increasing steps of 100 μA were tested in the first animal. However, we subsequently determined that stimulation intensities of 40, 140, 240 and 500 μA provided a good range for activating the circuits which were then selected for the experimental mapping of effective connectivity in both monkeys (Fig. 3). During a typical stimulation protocol, single pulses or bursts were repeated every 2 s for a total number of 40 trials (Fig. 2a). We used microstimulation to determine whether ACC and IPFC show effective connectivity. In half of the sessions, ACC neurons were stimulated and activation of the IPFC was recorded. If the recorded IPFC sites received input from ACC, then an evoked field potential (EFP) could be reliably observed for each ACC stimulation trial (e.g. Fig. 2b). In the other half of the sessions, we tested the directionality of the circuit by stimulating IPFC and recording from ACC in either the same or similar locations used for stimulation and recording in the preceding sessions. The depths of the stimulation and recording microelectrodes were adjusted under electrophysiological monitoring of the LFPs until EFPs were triggered. Data collection started 5 s before stimulation (pre-stimulation period) and continued for 5 s after stimulation (post-stimulation period). The 40 stimulations at the same stimulation and recording depths and locations were averaged for later analysis.

**Figure 1.**
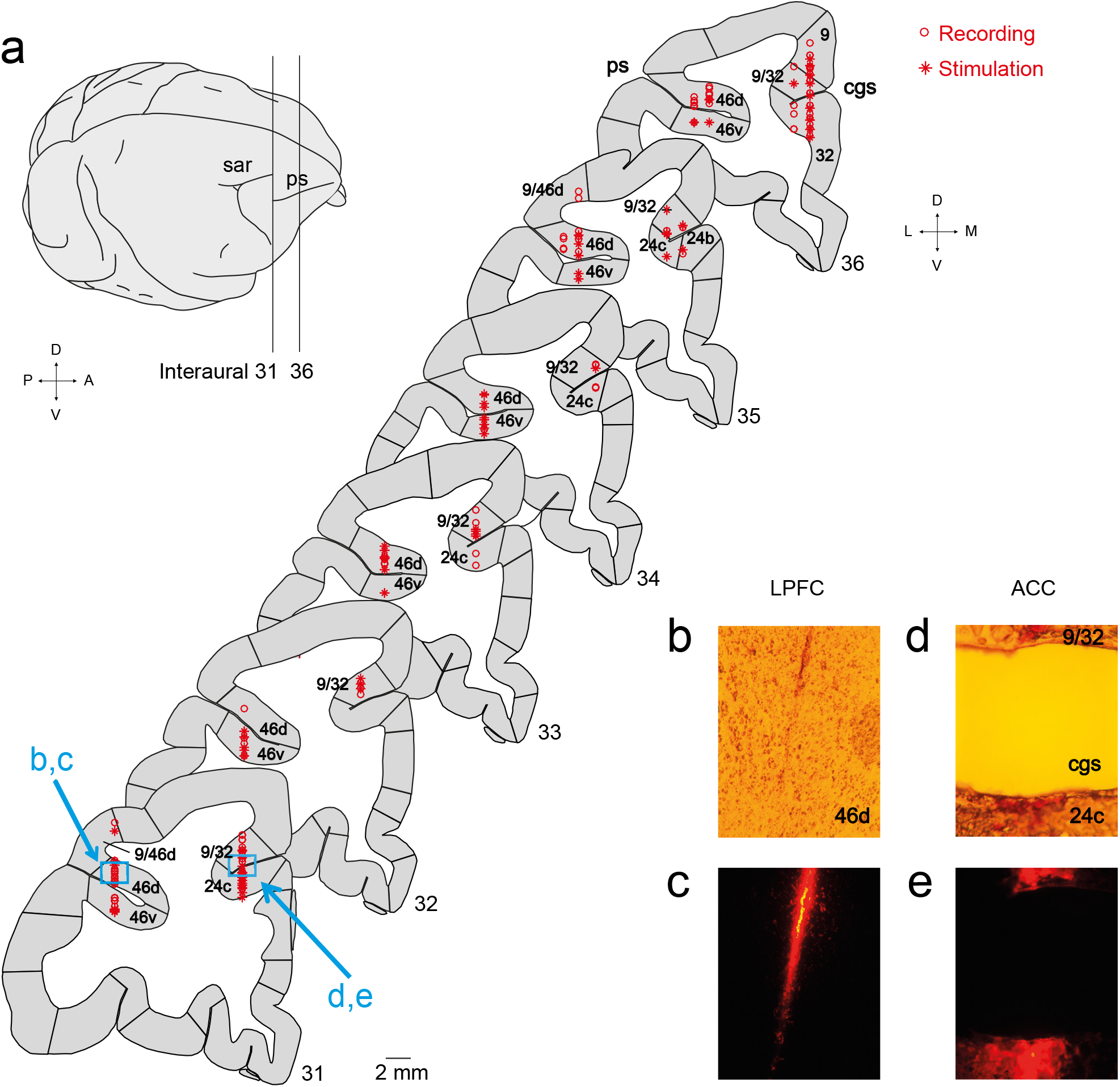
Anatomical reconstruction of the stimulation (open circles) and recording sites (asterisks) in ACC and IPFC areas. **a,** Right lateral surface of the rhesus monkey brain. The vertical lines depict the limits of the location of six plates in the coronal plane shown on the right. In each coronal plate the number at the bottom right shows the anterior-posterior distance relative to the interaural line (Paxinos et al., 2008). **b, d,** Bright-field micrograph of regions indicated in **a** (blue squares) for IPFC area 46d (**b**) and ACC areas 9/32 and 24c (**d**). **c, e,** Fluorescent photomicrographs of the same fields and sections in **b** and **d** showing the dye traces left by microelectrodes. The punctuate areas of high fluorescence are cell bodies that have incorporated the dye (DiI). Abbreviations: sar, superior arcuate sulcus; ps, principal sulcus; cgs, cingulate sulcus; IPFC, lateral prefrontal cortex; ACC, anterior cingulate cortex. Axes: D, dorsal; V, ventral; A, anterior; P, posterior; L, lateral; M, medial.

**Figure 2.**
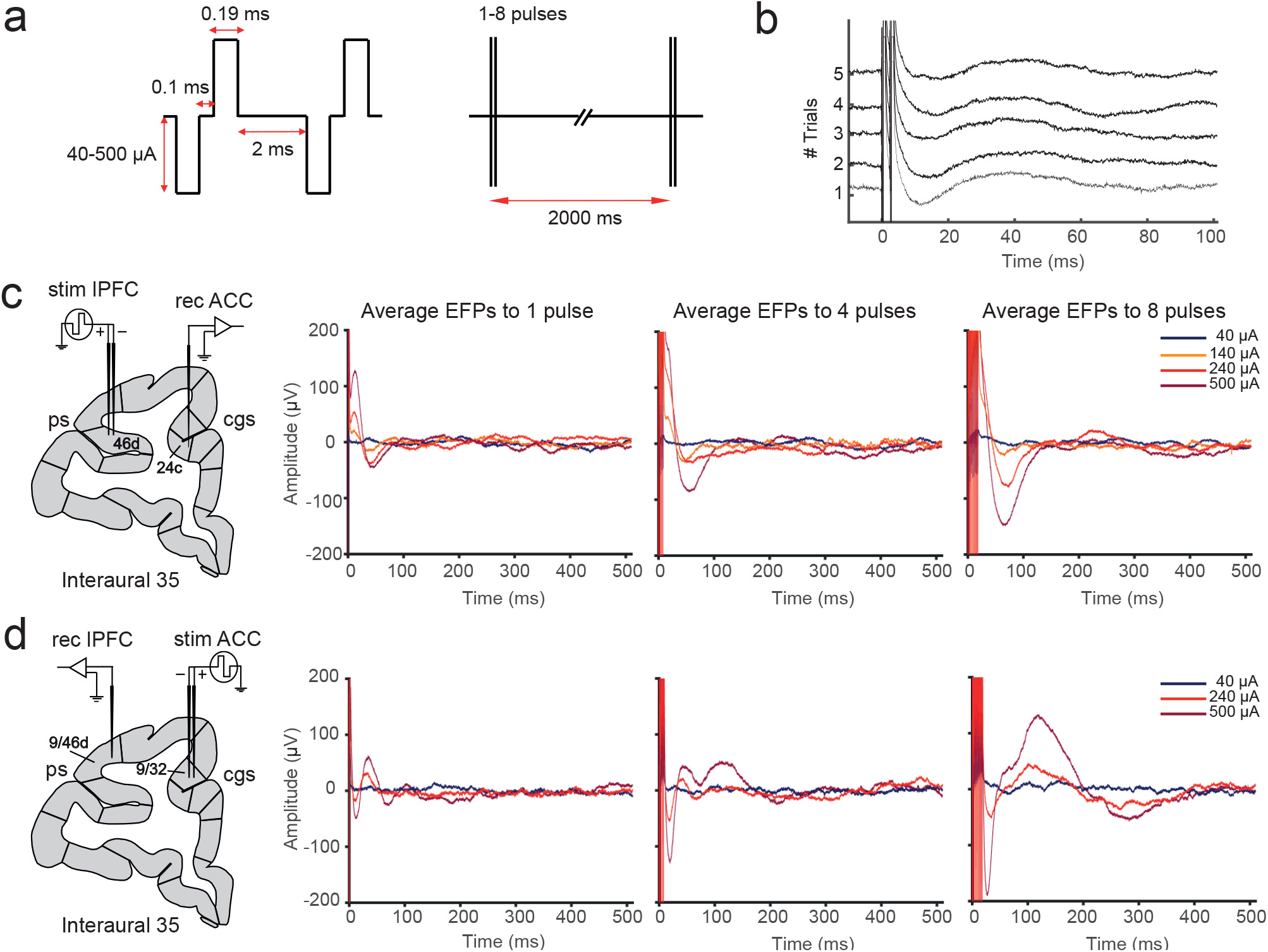
Example procedures for stimulating and recording evoked field potentials (EFPs). **a,** Parameters of the biphasic pulses used for stimulation. **b,** Example wideband IPFC recording during 5 example stimulations of 2 pulses bursts. Vertical lines correspond to the stimulation artifacts. Reliable and consistently activated EFPs follow each stimulation burst. **c,** Left, example experimental setup for IPFC stimulation and ACC recording. Right, EFP averages over 40 trials of stimulation recorded in ACC. The amplitude of evoked potentials increased from 1 to 8 pulses and 140 to 500 μA stimulation intensity. Note that no EFPs were elicited with 40 μA stimulation current. **d,** Same description as in **c**, but for examples of EFPs recorded in IPFC when ACC was stimulated.

**Figure 3.**
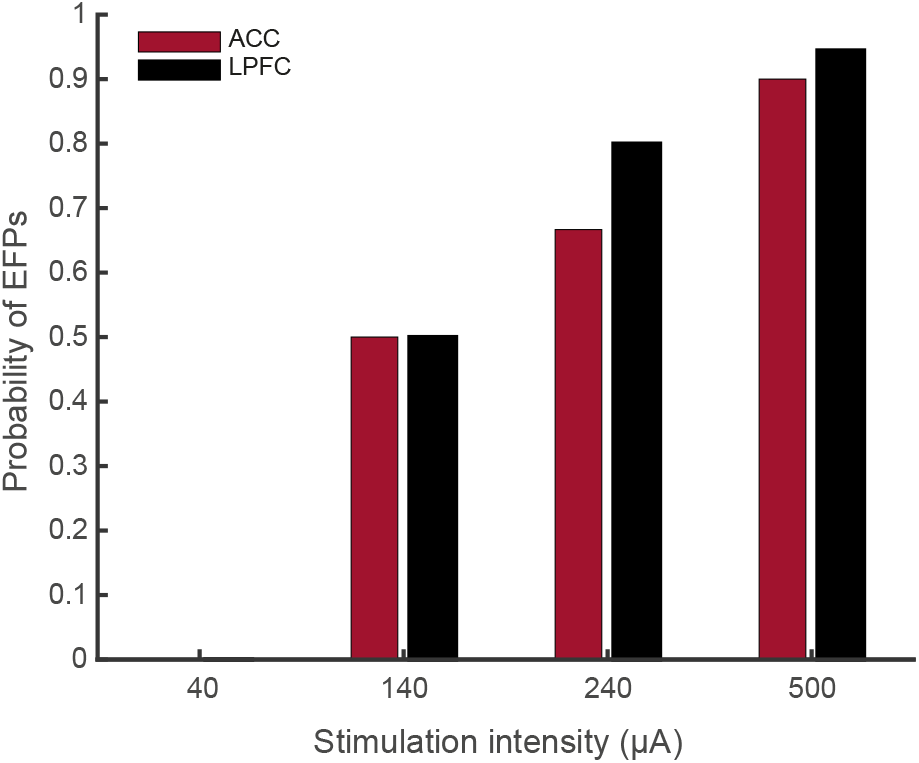
Probability of eliciting long-distance EFPs across sites. Each bar represents the probability of causing an EFP as function of the stimulation amplitude. Following single pulse stimulation, the least stimulation intensity which produced an EFP in 50 % of the recording sites in ACC and IPFC areas is 140 μA.

### Data analysis

Responses were analyzed offline using matlab (The Mathworks, Inc.). The majority of EFPs recorded in ACC and IPFC showed a multiphasic pattern with 2-3 points of inflection (Fig. 4). The EFPs were averaged and points of inflection were detected using a semi-automatic peak detection procedure using custom matlab scripts. First, peaks were detected automatically. Then, each detected peak was validated under visual guidance. We also utilized the first and second derivatives of the LFP signals following stimulation as additional visual confirmation of the precise time of peaks. For each peak, we extracted the amplitude and latency of the components following the procedure and terminology in Wallace et al., (2014). Chi-square and paired *t* tests were performed to determine the statistical differences; a significant difference level was set at p < 0.05.

**Figure 4.**
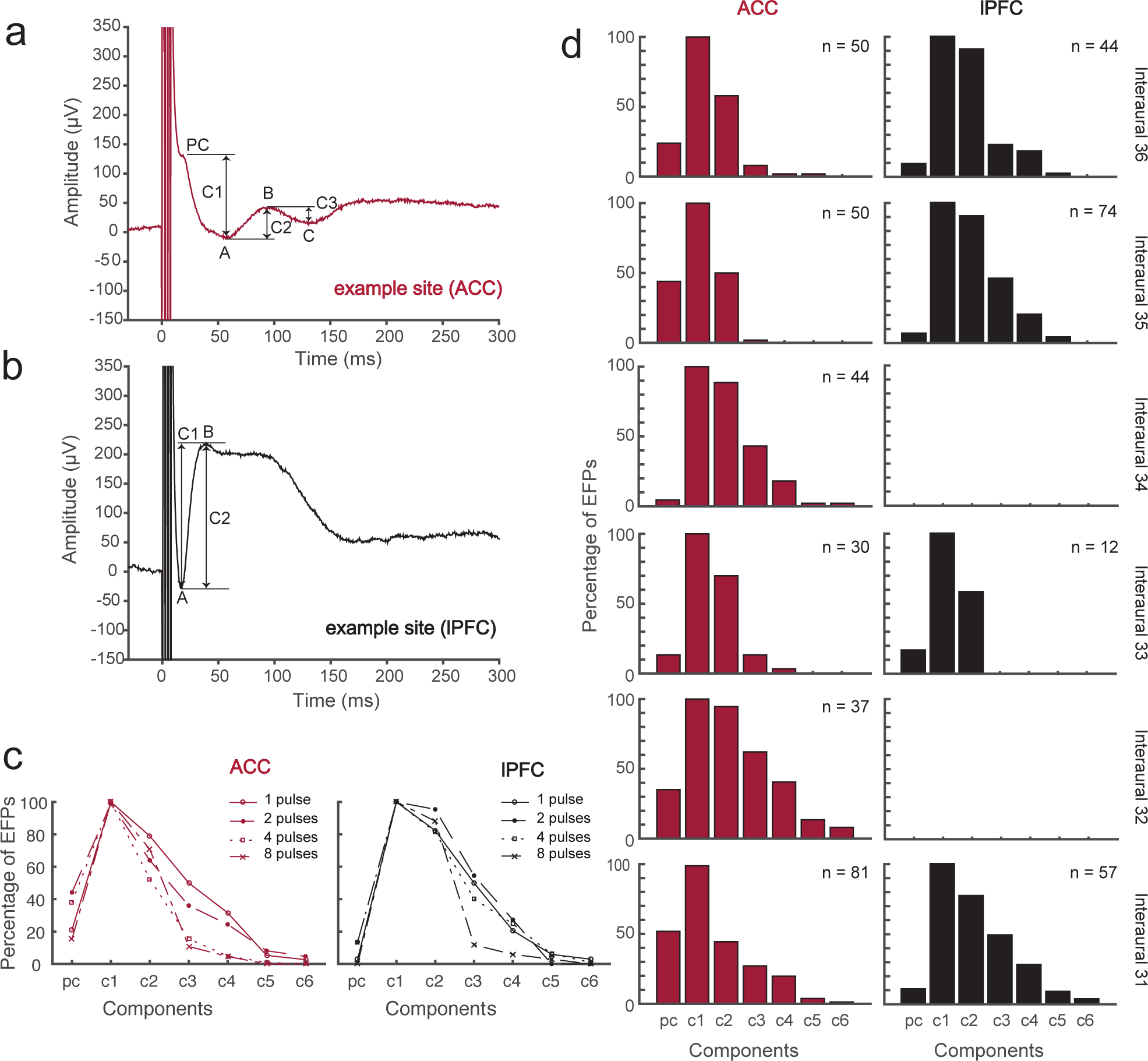
Typical averaged EFPs elicited in ACC (**a**) and IPFC (**b**). Vertical lines correspond to the stimulation artifacts. Stimulation parameters: 4 pulses burst, 500 μA. Letters mark the distinct points of inflection and arrows show how these were converted into components PC and C1-C3 (Wallace et al., 2014). **c**, Percentage of EFPs (*y-axis*) showing the individual components (*x-axis*) when stimulated with different number of pulses. **d,** Histograms showing the distribution of ACC and IPFC waveform components across the different interaural coordinates. EFPs recorded during mapping experiments are not included. The IPFC circuits showed significantly more periods (C2 to C4 components) of the evoked field potential (*see* text for details). We could not find evoked responses in IPFC for coordinates 34 and 32, and found very sparse responses in coordinate 33. Note the multiphasic property of the EFPs.

### Histology

At the end of the experiments, we confirmed the recording and stimulation sites in the first monkey by marking microelectrode tracks with a fluorescent dye (DiCarlo et al., 1996). Microelectrodes were coated with a commercial dye (DiI, 50 mg/ml, Sigma-Aldrich) before penetrating them at specific experimental stereotaxic coordinates in IPFC and ACC. Coating the microelectrodes with Dil allows each penetration to be located because it has a unique fluorescent absorption/emission signatures (550/565 nm peak) observable using a fluorescent microscope (Fig. 1c, e). The technique involves no treatment of the tissue other than standard perfusion and sectioning. The monkey was deeply anesthetized and perfused through the heart with saline followed by 3 % formaldehyde. Each microelectrode penetration was marked on the surface map of the brain area. The brain was removed and kept in 3 % formaldehyde and cryo-protected with 20 % sucrose solution. Blocks of the brain were removed from the right hemisphere and sectioned into 50 μm-thick coronal slices parallel to the electrode tracks. Slices were collected into cold phosphate buffer and mounted on gelatin-coated slides. A fluorescent microscope was used to confirm the trajectories of the microelectrodes. Next, the slices were stained with cresyl violet to obtain anatomical information and stored.

## Results

We combined electrical microstimulation in one brain area and multi-electrode recordings in the other brain area to investigate the nature and pattern of neural activity that effectively links the ACC and IPFC in two rhesus macaque monkeys. We will first report the EFP responses using different microstimulation protocols, and then report the temporal profile, distance and depth dependence of stimulation triggered EFPs in ACC and IPFC.

### Stimulation Triggered Evoked Field Potentials and Their Temporal Response Profile

A total of 243 pairs of stimulating and recording sites were tested in two monkeys (Monkey S = 93; Monkey H = 150, Fig. 1a) under several stimulation protocols combining different number of pulses (1, 2, 4 and 8) and stimulation intensities (40, 140, 240 and 500 μA). Across all sessions 812 electrical stimulation protocols were tested. 479 protocols were measured to characterize the typical evoked responses using all pulse types, while 333 protocols were measured to map the depth profile of the microstimulation effects using only the effective protocols eliciting reliable evoked fields.

At the beginning of each experiment we mapped the EFP amplitudes for different stimulation intensities and number of stimulation pulses (Fig. 2c, d). Both in ACC and IPFC, EFPs were reliably evoked at a minimum stimulation intensity of 140 μA with an increased proportion of sites showing evoked fields with larger amplitudes (Fig. 3). Following the stimulation artifact, the majority of EFPs were multiphasic, showing up to six points of inflection (for example sites from the ACC and IPFC, see Fig. 4a,b). The first point was a low amplitude negative deflection, similar to a component that previous studies considered to be a putative presynaptic component (PC) because it was unaffected by glutamate blockade (Wallace et al., 2014). The PC was not always present and the probability of its occurrence was higher in EFPs recorded in ACC after IPFC stimulation (χ^2^ (1) = 24.61, p < 0.0001), following bursts of 2 and 4 pulses (Fig 4c, d). The majority of EFPs had an initial trough (*labeled ‘A’* in Fig. 4a, b), followed by a peak (‘B’) and a trough (‘C’).

The average duration between evoked components ranged from 30-60 ms, indicative of a ~16-30 Hz beta frequency band resonance (Fig. 5). The duration of these resonance cycles was shortest for initial C1→C2 evoked components than for later components in both ACC and IPFC (Fig. 5). Notably, the average cycle durations were longer in ACC than in IPFC (χ^2^ (1) = 20.88, p < 0.0001). When testing individual cycle durations for ACC versus IPFC we found significantly different durations for C1→C2 (Wilcoxon ranksum, 1683, p < 0.0001), and C3→C4 (Wilcoxon ranksum, 552, p < 0.005). There was no apparent difference in ACC versus IPFC for the durations of C2→C3 (Wilcoxon ranksum, 1717, p = 0.0992), and C4→C5 (Wilcoxon ranksum, 26, p = 0.9657).

**Figure 5.**
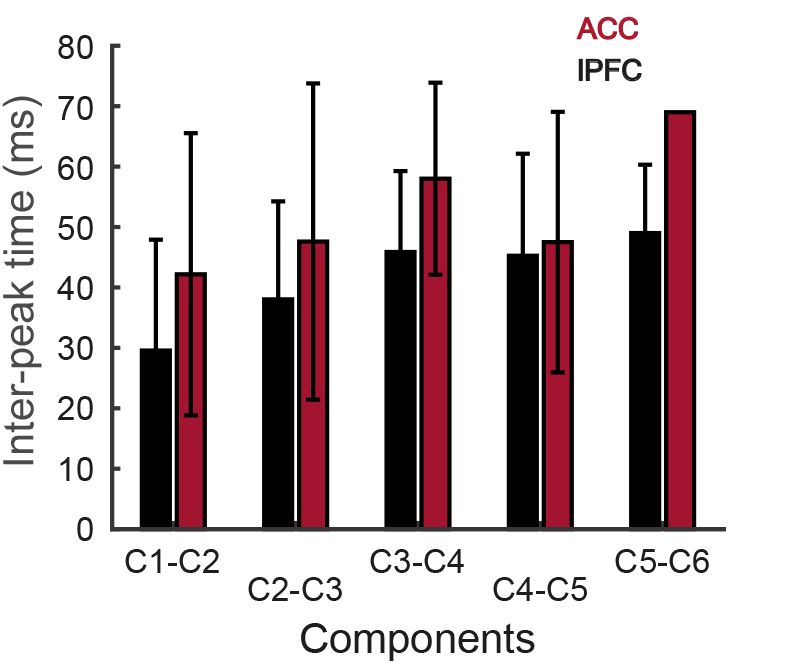
Average duration ± s.e.m between subsequent components of the multiphasic evoked responses in ACC and IPFC. We only had one ACC datapoint for the C5→C6 period.

Across stimulation – recording pairs we found differences in the number of components as a function of the number of pulses delivered during the stimulation (Fig. 4c). In general, the number of components for a given EFP decreased when using more stimulation pulses. In both areas a large proportion of EFPs following 8 pulses of burst stimulation had only 2 components (C1 and C2). However, the EFPs following single or double pulses showed a more complex waveform with 4 or more components.

### Anatomical Distribution of Stimulation Triggered Field Potentials

We next quantified the distribution of the PC and C1-C6 components at different interaural (anterior-to-posterior) coordinates (Fig. 4d). In IPFC, the number of elicited EFPs was significantly higher for the more anterior (interaural 36 and 35) and the most posterior (interaural 31) coordinates (t (4) = 5.68, p < 0.005). In contrast, for ACC sites the amount of elicited EFPs recorded was similar across all the interaural coordinates explored, although a slightly higher number of evoked fields were found at interaural coordinates 35, 36 and 31, but these did not reach statistical significance (t (4) = 2.10, p = 0.0517). For these locations, there were differences in the percentage of EFPs with PC, with higher percentage of PC’s in ACC (χ^2^ (1) = 42.38, p < 0.0001). The percentage of elicited C1 components in both areas was similar, but there were differences in the percentage of EFPs that showed C2-C4, with larger percentages for IPFC (C2: χ^2^ (1) = 14.70, p < 0.0001; C3: (χ^2^ (1) = 32.11, p < 0.0001); C4: (χ^2^ (1) = 18.98, p < 0.0001). Importantly, no EFPs were elicited in IPFC when stimulation in ACC was delivered at interaural coordinates 34 or 32, indicating that the anatomical connections between ACC and IPFC are not completely bidirectional at this point, at least with the explored stimulation-recording sites (Fig. 1). In fact, the number of EFPs recorded in both ACC and IPFC areas at interaural 33 was the lowest, compared with the other explored coordinates. For the IPFC, few EFPs were activated meaning a weaker connection in the direction of ACC to IPFC.

The overall relationship of stimulation intensity, number of stimulation pulses and EFP amplitude is shown in Fig. 6. For single and dual stimulation pulses there was a systematic increase in the mean EFP amplitude with increasing pulse strength in both the ACC (Fig. 6a) and IPFC (Fig. 6b). Stimulating with ≥ 3 stimulation pulses resulted in more variable relationship between stimulation intensity to EFP amplitudes, with reduced EFP amplitudes observed for most stimulation protocols involving 8 pulses.

**Figure 6.**
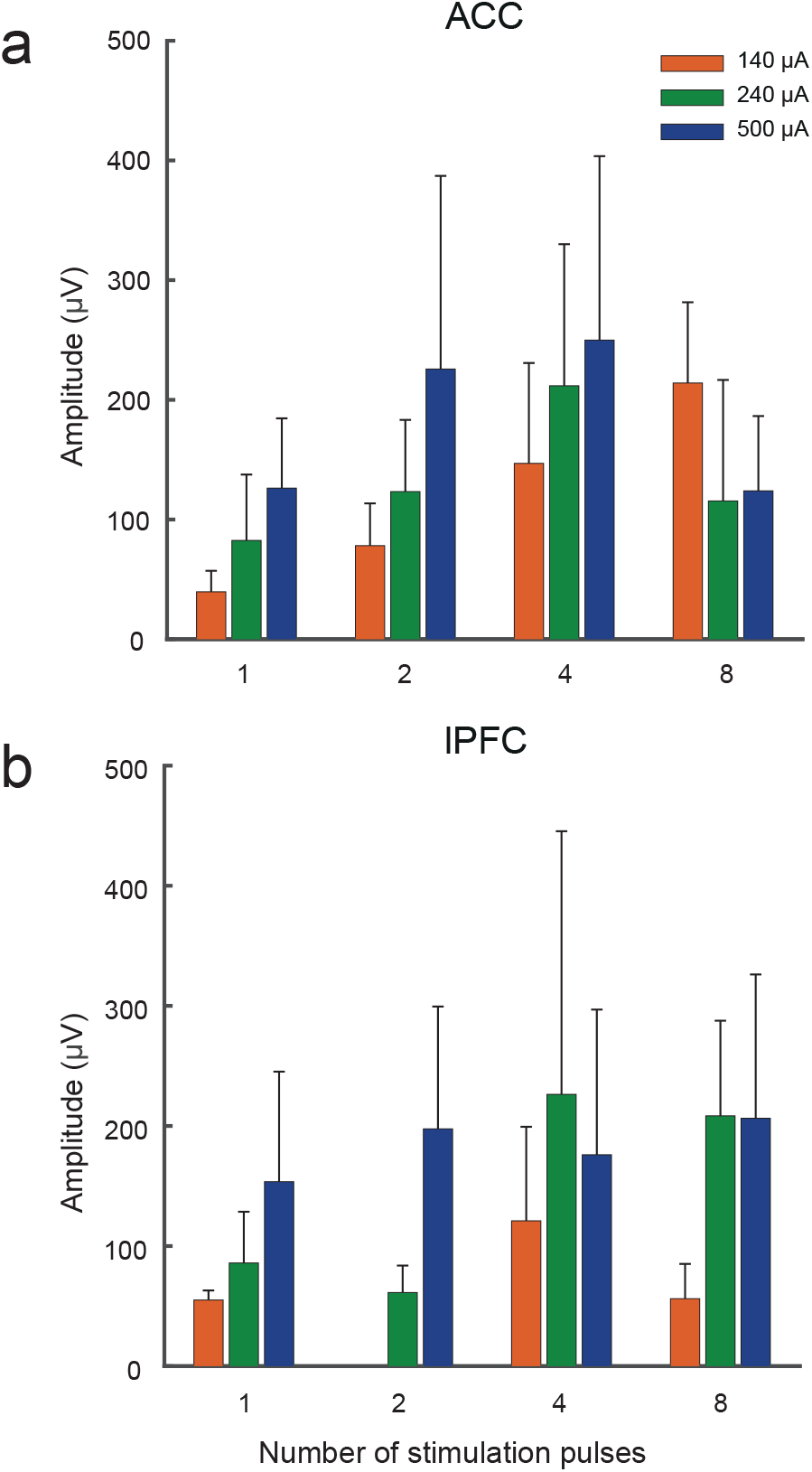
Relationship between stimulation intensity, number of pulses and EFP amplitude. **a,** ACC EFP amplitude modulation. **b,** IPFC EFP amplitude modulation. Each bar represents mean ± s.e.m of the C1.

### Latencies of Stimulation Triggered Evoked Field Potentials

The median peak latency of the C1 of the EFPs varied across the interaural coordinates and with increasing number of stimulation pulses (Fig. 7). Latencies for C1 were shortest when ACC was stimulated and IPFC is recorded from the most anterior (interaural 36 and 35) and posterior (interaural 31) coordinates. Also, the difference between these latency values increased with the number of pulses for these specific coordinates. Single pulse (IPFC: 10-14 ms; ACC: 28-38 ms), 2 pulse bursts (IPFC: 10-20 ms; ACC: 33-45 ms), 4 pulse bursts (IPFC: 18-20 ms; ACC: 45-57 ms) and 8 pulse bursts (IPFC: 24-59 ms; ACC: 70-79 ms). Interaural 33 was exceptional in that latencies were long and similar for both areas. The asymmetries in latencies found for the most anterior and posterior coordinates could indicate differential synaptic connectivity profiles in both directions.

**Figure 7.**
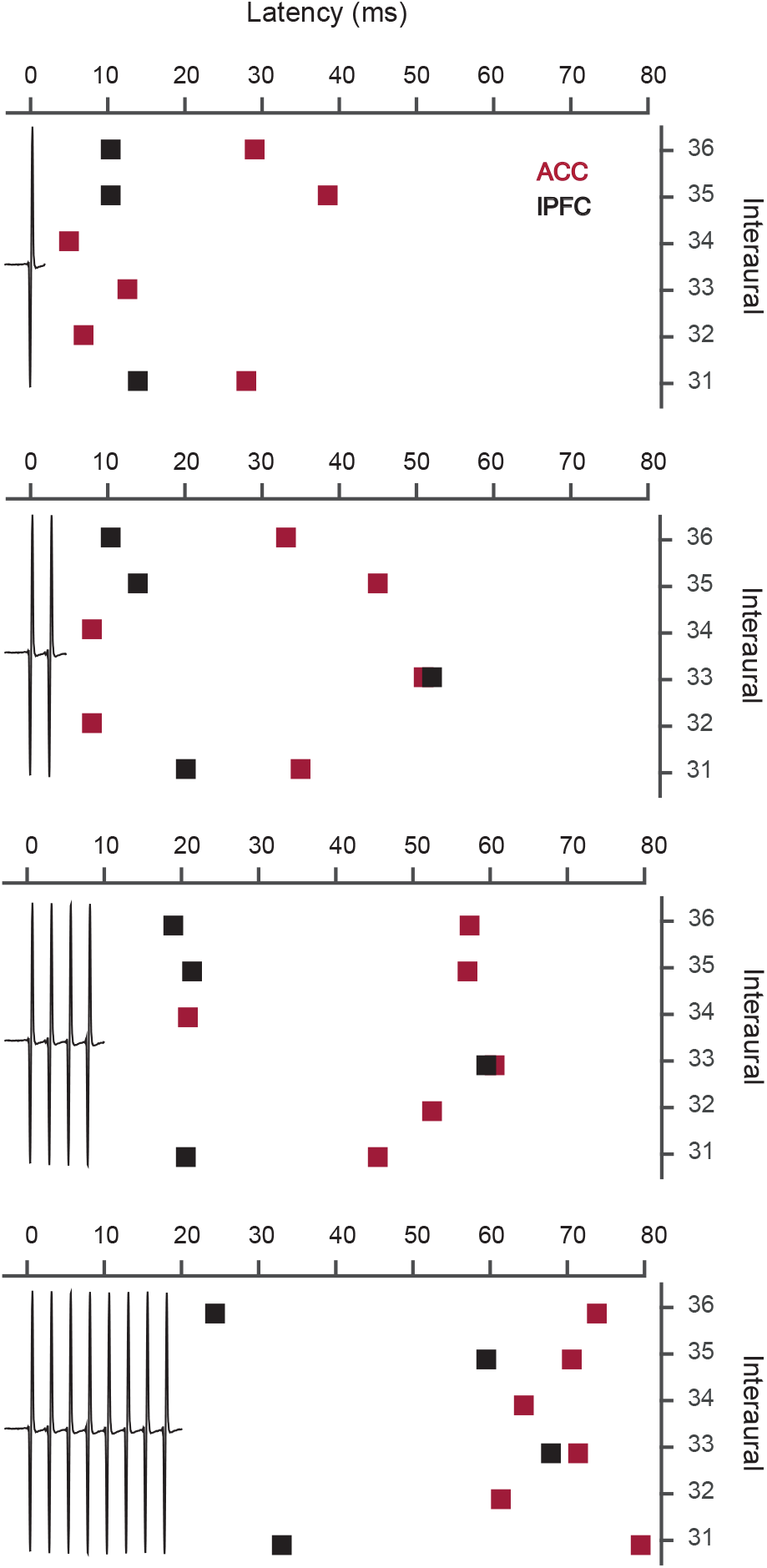
Latencies of EFPs at different interaural coordinates for 1-8 stimulation pulses. Shown are the median latencies of the C1 component of the EFPs in the ACC (dark red squares) and in the IPFC (black squares) at the different interaural coordinates (*y-axis*). The number of pulses is represented by the stimulation artifacts plotted at the actual trial time.

### Effects of Variation in Stimulation Depths on Evoked Field Potentials

We next explored whether long-distance microstimulation effects varied across laminar depth and were restricted to connections originating in grey matter. For this purpose, we mapped effective connectivity between ACC and IPFC at different depths across the cortex using stimulation protocols (2 or 4 pulse bursts at 500 μA supra-threshold intensity) which elicited evoked fields for most connections in the previous experiments (Fig. 3). Figure 8 shows the effects of changing the depths of stimulation microelectrodes on the EFPs in different ACC and IFPC subfields. During a representative mapping experiment shown in Fig. 8a (data from Monkey S), the recording microelectrodes remained at the same location in ACC (area 24c) while the stimulation microelectrodes was lowered into IPFC area 46d over a range of ~2mm until we observed a clear drop in the long-range evoked field amplitude. Across this depth range microstimulation had initially no effect on the ACC recording sites and then showed an ‘M’ like laminar effective connectivity profile for both ACC recording sites with increasing depth of the stimulation site. The dip in the middle of the depth profile spanned a ~600 μm window where the evoked field amplitudes were below the peak amplitudes at more superficial or deeper recording sites. In other mapping experiments with varying stimulation depth in the ACC and recordings in dorsal IPFC (Fig. 8b) or with varying stimulation depth in the ventral bank of the principle sulcus of the IPFC and recordings in ACC, we observed bell shaped depth profiles (Fig. 8b) or mixtures of bell shaped and M-shaped (Fig. 8c) depth profiles spanning ~1.5mm cortical distance in which stimulation triggered evoked fields. These findings indicate that systematic depth variations of effective connectivity might exist, and that overall, effective microstimulation effects at the intensities tested are restricted to cortical grey matter.

**Figure 8.**
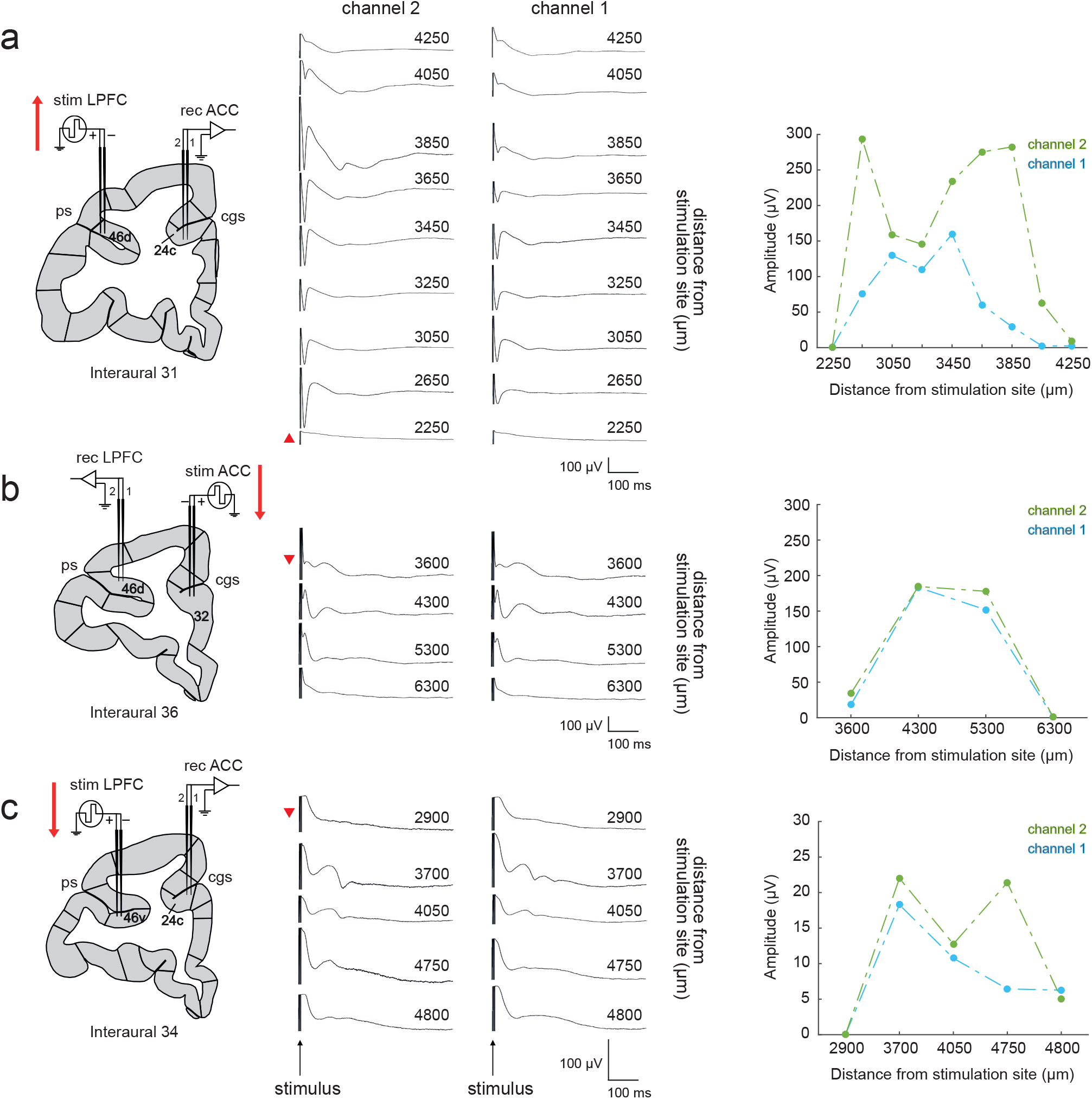
Representative inter-laminar mapping of the long-distance microstimulation effects between ACC and IPFC areas. **a**, Effects of changing the depth of the stimulation microelectrodes. Each EFP was recorded at the same depth in right area 24c (channels 1 and 2), with the position of the stimulation electrodes in IPFC area 46d changing in the ventrodorsal direction (red arrow represents de direction of the mapping). The red triangle to left of a specific recording trace depicts the starting mapping location and its direction. Numbers above each recording trace denote the distance of the recording from the stimulation site in μm. Stimulation parameters: 2 pulses burst, 500 μA. Plot on the right shows the layer-specific EFP amplitude modulation for the C1 and the extent of the microstimulation effects. **b**, Effects on IPFC area 46d EFPs of changing the depth of the stimulation electrodes in area 32. Stimulation parameters: 4 pulses burst, 500 μA. **c**, Effects on area 24c EFPs of changing the depth of the stimulation electrodes in area 46v. Stimulation parameters: 4 pulses burst, 500 μA. Abbreviations: ps, principal sulcus; cgs, cingulate sulcus.

## Discussion

We found a microelectrical stimulation protocol for mapping the strength, latency and temporal patterning of effective connectivity between the ACC and IPFC in the awake non-human primate. Effective synaptic connectivity was reliably evoked with 140 μA pulses in half of all connections in both directions, increasing systematically with increasing amplitudes up to ~500 μA. Whether two sites within the ACC and IPFC were connected depended on the precise anterior to posterior locations in a patchy way with no apparent linear gradient. Despite this heterogeneity, we observed reliable directional asymmetries of the ACC-IPFC connectivity. ACC stimulation triggered faster latency evoked components in the IPFC and resulted, more likely, in a multiphasic response pattern with 2-4 inflection points when compared to stimulating the same connection from the IPFC and measuring the evoked component in the ACC. These findings provide to the best of our knowledge the first delineation of effective connectivity among ACC and IPFC in the awake monkey brain, pointing to anatomically patchy connectivity that shows an asymmetric temporal profile. We speculate that these findings at the level of effective connectivity reflect patchy and asymmetric synaptic connectivity at the anatomical level, and might translate into differences in neuronal information flow during functional states that depend on ACC-IPFC contributions.

### Eliciting long-range connectivity with single and dual electrical pulses

Our study identified a bipolar, biphasic, constant-current electrical stimulation protocol that reliably evokes long-range LFP responses for most ACC-to-IPFC and IPFC-to-ACC long-range connections. This protocol deviates significantly from previous studies reporting long-range electrical stimulation effects on blood flow responses using functional magnetic resonance imaging (fMRI) (Matsui et al., 2011; Tolias et al., 2005; Ekstrom et al, 2008; Moeller et al., 2008; Field et al., 2008; Premereur et al., 2015; Logothetis et al., 2012). When testing protocols that are similar to those used in fMRI studies such as e.g. using trains of biphasic pulses (25-50 pulses) at 100 Hz with current intensities ranging from 20-240 *μ*A, we failed to observe reliable evoked fields. In our setting, it was sufficient to use single and dual pulse stimulation to trigger evoked fields at 140-500 *μ*A.

### Long-distance activation is multiphasic and could indicate circuit specific resonance

We found that a main characteristic of long-range evoked responses was a multiphasic temporal response pattern. For more than half of all connections microstimulation triggered a dynamic, multiphasic response with 2 or more peaks. These multiphasic EFPs displayed a recurrent negative-positive pattern of several cycles with decreasing amplitude suggestive of a recurrent excitatory loop (Wallace et al., 2014). This could indicate that neurons closer to the recording site in the activated long-distance area, either ACC or IPFC, respond more effectively to inputs of a particular low frequency range which is reflected in the shape of the EFP. We interpret this as resonance phenomenon that could help to understand the underlying mechanisms of synaptic integration (Buzsaki et al., 2012; Hutcheon and Yarom, 2000). Consistent with this interpretation, modeling studies suggest that resonance, as measured in LFP signals, is well retained close to the cell somata when the synaptic inputs are concentrated at the apical dendrites (Ness et al., 2016). Resonance phenomena are more difficult to measure when inputs are distributed more uniformly across apical and basal dendritic trees (Ness et al., 2016). We believe that the widespread rhythmic resonance effects we observed even following single, non-rhythmic input pulses makes it likely that a similar resonance phenomenon might be effective during intrinsic information processing, having possibly widespread consequences as outlined in the framework of neuronal ‘communication though resonance’ (CTR) (Hahn et al., 2014). CTR is built on the observation that in local circuits with asynchronous-irregular firing activity without apparent ongoing oscillations in the LFP activity, already subtle resonance can lead to the amplification of input over successive cycles within the resonant frequency (Hahn et al., 2014). According to these computational insights the observed beta band resonance in both ACC and LPFC circuits, could reflect an intrinsic mechanism for enhanced recurrent communication between both areas triggered by task demands or internal state changes.

One important contribution to the resonance of a circuit are the time constants of inhibitory synapses that influence the period of rhythmic responses (Womelsdorf, Valiante et al., 2014). Our finding that the IPFC circuits showed significantly more periods (C2 to C4 components) of the evoked field potential might therefore be due to a strong pool of inhibitory synapses. Consistent with this interpretation, a previous anatomical study found that the population of parvalbumin expressing (PV+) inhibitory interneurons was more than twice as numerous in IPFC compared with ACC, while slower acting calbindin (CB+) expressing interneurons were more numerous in ACC than in IPFC (Dombrowski et al., 2001). An alternative interpretation posits that due to a lower density of pyramidal neurons in the ACC compared to IPFC as well as the higher frequency, amplitude and duration of IPSCs within the ACC relative to the IPFC, the ACC robustly and rapidly inhibited the excitatory input (Medalla et al., 2017). These different profiles of inhibition in IPFC and ACC might contribute to the different temporal patterning of the evoked response. We believe that future modeling studies could clarify the relationship of local resonance and the composition of the resonating neural populations.

### Effective connectivity shows discontinuities similar to anatomical connectivity

By varying the anterior-to-posterior location of stimulation and recording sites we were able to discern that the strength and latencies of electrically evoked field components could vary abruptly from one anterior-to-posterior section to the next section 1mm away (Fig. 4 and Fig. 6). We believe that such discontinuities in the strength of effective synaptic connectivity might reflect discontinuities of the underlying anatomical connectivity. Previous anatomical tracing studies have shown that both brain regions we recorded and stimulated, rostral ACC and the ventral and dorsal bank of the principal sulcus (IPFC), are reciprocally connected over a wide anterior to posterior range (Arikuni et al., 1994; Lu et al., 1994). However, visually inspecting the density and distribution of cells in the IPFC that were retrogradely labeled by large tracer injections in the ACC reveals dramatic variations from one coronal slice to the next (Arikuni et al., 1994). While cells can show labeling over several coronal slices, occasionally a slice shows reduced labeling or lacks labeling altogether. These anatomical discontinuities could well underlie the abrupt variations in latencies we observed between interaural 31 mm and 32 mm or between interaural 34 mm and 35 mm (Fig. 5 upper panels).

A similar anatomical origin might also underlie the two different depth profiles of effective connectivity we observed. We found that long-range evoked fields showed bell-shaped or ‘M’-shaped profiles when lowering the stimulation electrodes through the layers inside the grey matter (Fig. 7). These two types of patterns can likewise be seen in the density profiles of labelled projection cells traced with retrograde tracers between ACC and IPFC. For example, Arikuni et al. (1994) presented a case with tracer injection in the rostral ACC that caused patchy, retrograde labeling of cells in the ventral bank of the principal sulcus (IPFC) in the form of a superficial cluster and a deep layer cluster similar to an M-pattern (e.g. Fig. 2, sections 14 and 16 in Arikuni et al. 1994). Similar patterns with two layers of retrogradely labeled cells are evident in other studies (e.g. Fig. 9c, section 2 in Morecraft et al., 2012), suggesting that cortico-cortical connections between ACC and IPFC proceed through two separate routes in upper and lower layers. In addition to the ‘two striped’, or ‘M-shaped’ pattern in anatomically tracing, several slices show a more homogeneous labeling with only a single cluster of cells across laminae with the highest density of connections in the center of the cluster, reminiscent of a bell-shaped profile. These anatomical studies thus provide connectivity profiles similar to those that we observed in profiles of effective connectivity in the awake monkey.

### Latency differences of evoked fields suggest asymmetric ACC-IPFC effective connectivity

We found that the latencies of EFPs recorded from IPFC when ACC was stimulated was systematically shorter, across the different number of stimulation pulses, than the latency in ACC when IPFC was stimulated at interaural coordinates 36, 35 and 31. This asymmetry could indicate differential synaptic connectivity profiles. Medalla and Barbas (2010) addressed the issue of the mediolateral prefrontal activity at the synaptic level. The pathways from area 32 to layers I-III of dorsolateral areas 10, 46 and 9 had in common a higher prevalence of large boutons innervating putative inhibitory neurons. Their functional interpretation is that ACC pathways may help reduce excessive noise in IPFC during demanding cognitive tasks by strongly activating inhibitory neurons through large terminals. The size of boutons is positively correlated with the number of synaptic vesicles and the probability of release with each action potential, features that increase the synaptic efficacy and, presumably shorten the latency of the transmitted signals (Medalla et al., 2017).

In addition to this synaptic explanation, the asymmetries in latencies might also relate to an overall lower cell density in ACC compared with the IPFC superficial layers II and III (Dombrowski et al., 2001; Gabbott and Bacon 1996). A larger cell density will allow a faster spread of input current in the receiving circuit (McIntyre and Grill, 2000), and hence could also underlie faster evoked responses in the IPFC than in ACC. Although we cannot eliminate the possibility of more complex synaptic pathways between the stimulating and recording sites, the predictable nature of EFPs based on the electrode locations and their localization suggested that an organized, systematic flow of information between ACC and IPFC subfields exists.

### Conclusion

In summary, our study revealed systematic asymmetries in the latencies and temporal patterning of ACC-IPFC effective connectivity, suggesting a more rapid and more temporally patterning of ACC → IPFC effects than IPFC → ACC effects. These results identified a parameter range for the intensity and number of biphasic cathode-first-anode-second pulse types that reliably evoke field potential flow in the receiving brain circuit. This overall pattern of effective connectivity was complemented by anisotropies of stimulation effects across layers, and by patchy stimulation effects that showed stronger and weaker connectivity at anatomical sites spaced as little as 1mm away from each other. We believe that the findings of systematic asymmetries in effective connectivity might be reflected in asymmetries of functional connectivity and could originate from complex variations of synaptic connectivity between the ACC and IPFC. Future studies could clarify this suggestion by combining measuring effective and functional connectivity protocols in the same experimental set up and applying synaptic dual tracers for antero- and retrograde labeling of inter-area connections. Such a multi-method approach could resolve how structural and functional connectivity relate to each other to subserve the network level functions that co-activation of ACC and IPFC realizes during goal directed behavior.

## Compliance with ethical standards

### Ethical approval

All animal care and experimental procedures performed in this study have been approved by the local ethics committee, the York University Council on Animal Care, were in accordance with the Canadian Council on Animal Care guidelines, and are in agreement with the 1964 Helsinki declaration and its later amendments.

## Acknowledgments

This work was supported by a grant from the Canadian Institutes of Health Research (T.W.). CIHR Grant MOP_102482. The funders had no role in study design, data collection and analysis, the decision to publish, or the preparation of this manuscript. Authors would like to thank Hongying Wang for technical support.

## References

Alexander WH, Brown JW (2011) Medial prefrontal cortex as an action-outcome predictor. Nat Neurosci 14:1338–1344.

Arikuni T, Sako H, Murata A (1994) Ipsilateral connections of the anterior cingulate cortex with the frontal and medial temporal cortices in the macaque monkey. Neurosci Res 21:19–39.

Buzsaki G, Anastassiou CA, Koch C (2012) The origin of extracellular fields and currents—EEG, ECoG, LFP and spikes. Nat Rev Neurosci 13:407–420.

DiCarlo JJ, Lane JW, Hsiao SS, Johnson KO (1996) Marking microelectrode penetrations with fluorescent dyes. J Neurosci Methods 64:75–81.

Dombrowski, S.M., Hilgetag, C.C., and Barbas, H. (2001). Quantitative architecture distinguishes prefrontal cortical systems in the rhesus monkey. Cereb Cortex 11, 975–988.

Ekstrom LB, Roelfsema PR, Arsenault JT, Bonmassar G, Vanduffel W (2008) Bottom-up dependent gating of frontal signals in early visual cortex. Science 321:414–417.

Field CB, Johnston K, Gati JS, Menon RS, Everling S (2008) Connectivity of the primate superior colliculus mapped by concurrent microstimulation and event-related fMRI. PLoS One 3:e3928.

Gabbott, P.L., and Bacon, S.J. (1996). Local circuit neurons in the medial prefrontal cortex (areas 24a,b,c, 25 and 32) in the monkey: II. Quantitative areal and laminar distributions. The Journal of comparative neurology 364, 609–636.

Hahn, G., Bujan, A.F., Fregnac, Y., Aertsen, A., and Kumar, A. (2014). Communication through resonance in spiking neuronal networks. PLoS computational biology 10, e1003811.

Hayden, B.Y., Heilbronner, S.R., Pearson, J.M., and Platt, M.L. (2011). Surprise signals in anterior cingulate cortex: neuronal encoding of unsigned reward prediction errors driving adjustment in behavior. J Neurosci 31, 4178–4187.

Hutcheon B, Yarom Y (2000) Resonance, oscillation and the intrinsic frequency preferences for neurons. Trends Neurosci 23:216–222.

Kennerley SW, Walton ME, Behrens TE, Buckley MJ, Rushworth MF (2006) Optimal decision making and the anterior cingulate cortex. Nat Neurosci 9:940–947.

Logothetis NK, Eschenko O, Murayama Y. Augath M, Steudel T, Evrard HC, Besserve M, Oeltermann A (2012) Hippocampal-cortical interaction during periods of subcortical silence. Nature 491:547–553.

Lu MT, Preston JB, Strick PL (1994) Interconnections between the prefrontal cortex and the premotor areas in the frontal lobe. J Comp Neurol 341:375–392.

Matsui T, Tamura K, Koyano KW, Takeuchi D, Adachi Y, Osada T, Miyashita Y (2011) Direct comparison of spontaneous functional connectivity and effective connectivity measured by intracortical microstimulation: An fMRI study in macaque monkeys. Cereb Cortex 21:2348–2356.

McIntyre, C.C., and Grill, W.M. (2000). Selective microstimulation of central nervous system neurons. Ann Biomed Eng 28, 219–233.

Medalla M, Barbas H (2010) Anterior cingulate synapses in prefrontal areas 10 and 46 suggest differential influence in cognitive control. J Neurosci 30:16068–16081.

Moeller S, Freinwald WA, Tsao DY (2008) Patches with links: A unified system for processing faces in the macaque temporal lobe. Science 320:1355–1359.

Montgomery EB (2010) Deep brain stimulation programming: Principles and practice. Oxford University Press.

Morecraft RJ, Stilwell-Morecraft KS, Cipolloni PB, Ge J, McNeal DW, Pandya DN (2012) Cytoarchitecture and cortical connections of the anterior cingulate and adjacent somatomotor fields in the rhesus monkey. Brain Res Bull 87:457–4997.

Medalla M, Gilman JP, Wang JY, Luebke JI (2017) Strength and diversity of inhibitory signaling differentiates primate anterior cingulate from lateral prefrontal cortex. J Neurosci 37(18): 4717–4734.

Ness TV, Remme MWH, Einevoll GT (2016) Active subthreshold denditric conductances shape the local field potential. J Physiol 594:3809–3825.

Oemisch M, Westendorff S, Everling S, Womelsdorf T (2015) Interareal spike-train correlations of anterior cingulate and dorsal prefrontal cortex during attention shifts. J Neurosci 35:13076–13089.

Paxinos G, Huang XF, Petrides M, Toga AW (2008) The rhesus monkey brain in stereotaxic coordinates. Academic Press.

Premereur E, Van Dromme IC, Romero MC, Vanduffel W, Janssen P (2015). Effective connectivity of depth-structure-selective patches in the lateral bank of the macaque intraparietal sulcus. PLoS Biol 13:e1002072.

Rothe, M., Quilodran, R., Sallet, J., and Procyk, E. (2011). Coordination of high gamma activity in anterior cingulate and lateral prefrontal cortical areas during adaptation. The Journal of neuroscience: the official journal of the Society for Neuroscience 31, 11110–11117.

Rushworth MF, Noonan MP, Boorman ED, Walton ME, Behrens TE (2011) Frontal cortex and reward-guided learning and decision making. Neuron 70:1054–1069.

Shenhav, A., Cohen, J.D., and Botvinick, M.M. (2016). Dorsal anterior cingulate cortex and the value of control. Nat Neurosci 19, 1286–1291.

Tolias AS, Sultan F, Augath M, Oeltermann A, Tehovnik EJ, Schiller PH, Logothetis NK (2005) Mapping cortical activity elicited with electrical microstimulation using fMRI in the macaque. Neuron 48: 901–911.

Voloh B, Valiante TA, Everling S, Womelsdorf T (2015). Theta–gamma coordination between anterior cingulate and prefrontal cortex indexes correct attention shifts. Proc Natl Acad Sci U S A 112:8457–8462.

Voloh, B., and Womelsdorf, T. (2017). Cell-Type Specific Burst Firing Interacts with Theta and Beta Activity in Prefrontal Cortex During Attention States. Cereb Cortex, 1–17.

Wallace J, Jackson RK, Shotton TL, Munjal I, McQuade R, Gartside SE (2014) Characterization of electrically evoked field potenials in the medial prefrontal cortex and orbitofrontal cortex of the rat: Modulation by monoamines. Eur Neuropsychopharmacol 24:321–332.

Womelsdorf T, Ardid S, Everling S, Valiante TA (2014) Burst firing synchronizes prefrontal and anterior cingulate cortex during attentional control. Curr Biol 24:2613–2621.

Womelsdorf, T., Valiante, T.A., Sahin, N.T., Miller, K.J., and Tiesinga, P. (2014). Dynamic circuit motifs underlying rhythmic gain control, gating and integration. Nat Neurosci 17, 1031–1039.

Womelsdorf T, Everling S (2015) Long-range attention networks: circuit motifs underlying endogenously controlled stimulus selection. Trends Neurosci 38:682–700.

